# Distinct beta burst motifs exhibit opposing error relationships during motor adaptation

**DOI:** 10.64898/2026.03.06.710026

**Authors:** Quentin Moreau, Maciej J. Szul, Sébastien Daligaut, Denis Schwartz, James J. Bonaiuto

## Abstract

Beta-band activity (13-30 Hz) is a hallmark of human movement, yet a unifying account of its functional role remains unresolved. Although typically described as a sustained oscillation, beta activity is increasingly recognised to consist of transient bursts. More recently, beta bursts have been shown to exhibit heterogeneous waveforms. Here, we ask whether variability in burst shape corresponds to separable computational roles during motor adaptation. Using high-precision MEG, we recorded neural activity while participants performed a visuomotor rotation task under either implicit (sensorimotor adaptation) or explicit (strategic re-aiming) learning conditions. Conventional metrics, beta power and burst rate, showed context-dependent modulation during preparation but provided limited insight into trial-by-trial behaviour. In contrast, sorting bursts according to their waveforms revealed a repertoire of burst types with dissociable temporal dynamics and context-dependent modulation. Crucially, during post-movement evaluation, distinct burst subtypes showed opposing and temporally specific relationships with behavioural error: one subtype decreased with increasing error, whereas others increased. Together, these findings indicate that beta activity comprises separable transient events with specific computational roles, and that accounting for waveform diversity is essential for understanding how cortical beta supports adaptive behaviour.

## Introduction

Beta band activity (13-30 Hz) is reliably modulated during voluntary movement, showing a reduction prior to and during movement (movement-related beta decrease - MRBD), followed by a post-movement beta rebound (PMBR ^1–3)^. Despite decades of study, the functional significance of these modulations remains unresolved. Interpretations based solely on changes in beta power, ranging from cortical idling ^7^ to long-range communication ^5^ or integrative sensorimotor processing ^4^ have failed to converge on a unifying account.

Recent evidence suggests that beta activity is not a sustained oscillation, but is expressed as transient bursts at the single-trial level ^9–13^. Averaging across trials gives rise to the appearance of rhythmic activity in the frequency domain, whereas single-trial analyses reveal discrete burst events ^14–16^. Importantly, these bursts are not morphologically uniform ^7,8^ : waveform-resolved analyses indicate structured diversity beyond that indicated by time-frequency features, with different burst types showing distinct rate dynamics during motor behavior ^10,11,17–19^. These findings raise the possibility that beta activity comprises multiple functionally distinct transient motifs, rather than a unitary oscillatory process. However, whether specific burst types support different computational components of motor learning remain unclear.

Within motor learning, motor adaptation paradigms provide a principled framework to test this possibility because they implicate several distinct computational mechanisms with clear behavioral hallmarks^17^. Adaptation to visuomotor perturbations relies on at least two partially dissociable processes: implicit recalibration driven by sensorimotor prediction error, and explicit strategies including conscious re-aiming^18,19^. These processes exhibit distinct behavioral signatures, most notably the presence of aftereffects in implicit learning ^30,31^ and their absence in purely strategic adaptation. This dissociation provides a framework to test whether distinct waveform-defined burst types show differential relationships with behavior across implicit and explicit learning.

Here we classified beta bursts at the single-trial level according to their waveform features and examined their modulation during movement preparation and post-movement evaluation. Participants were assigned to either an implicit adaptation group exposed to a constant visuomotor perturbation, or an explicit group for whom trial-by-trial perturbations were cued predictively. We hypothesized that distinct waveform-defined burst types would show dissociable and temporally specific relationships with behavioral error, and that their expression during movement preparation would be modulated by learning context. We show that specific burst types exhibit differential temporal dynamics across movement phases, context-dependent modulation during learning, and distinct, often opposing relationships with movement error. These findings indicate that variability in beta burst waveforms is functionally meaningful, supporting a view of beta activity as comprising a structured repertoire of transient events with separable computational roles in motor adaptation.

## Results

### Behavior dissociates implicit and explicit motor learning

Participants were asked to perform ballistic joystick movements with their right hand to reach a visual target. Each trial began with presentation of a circular random dot kinematogram (RDK) with coherent motion in either a clockwise or counterclockwise direction at one of four coherence levels (no coherence, low, medium, or high). Five potential targets were positioned along the RDK’s edge, with one appearing larger and green, indicating the reach target. After a delay, the random dot kinematogram (RDK) disappeared, followed by the disappearance of the four non-target stimuli, which served as the go cue for participants to move a joystick-controlled cursor to the remaining target within 1 second (Figure 1A). Participants were randomly assigned to either an implicit (*N* = 18) or explicit (*N* = 20) group. All participants first completed a baseline block during which no visuomotor rotation was applied. This was followed by six adaptation blocks. In the implicit group, a constant −30° visuomotor rotation was applied during the reach (i.e. the visually presented cursor moved −30° relative to the movement of the joystick), irrespective of the direction of coherent motion of the random dot kinematogram (RDK). In contrast, the explicit group experienced a rotation that was cued by the direction of RDK coherent motion, enabling strategic re-aiming to compensate for it (Figure 1B). Finally, all participants completed a washout block during which there was no visuomotor perturbation.

**Figure 1.**
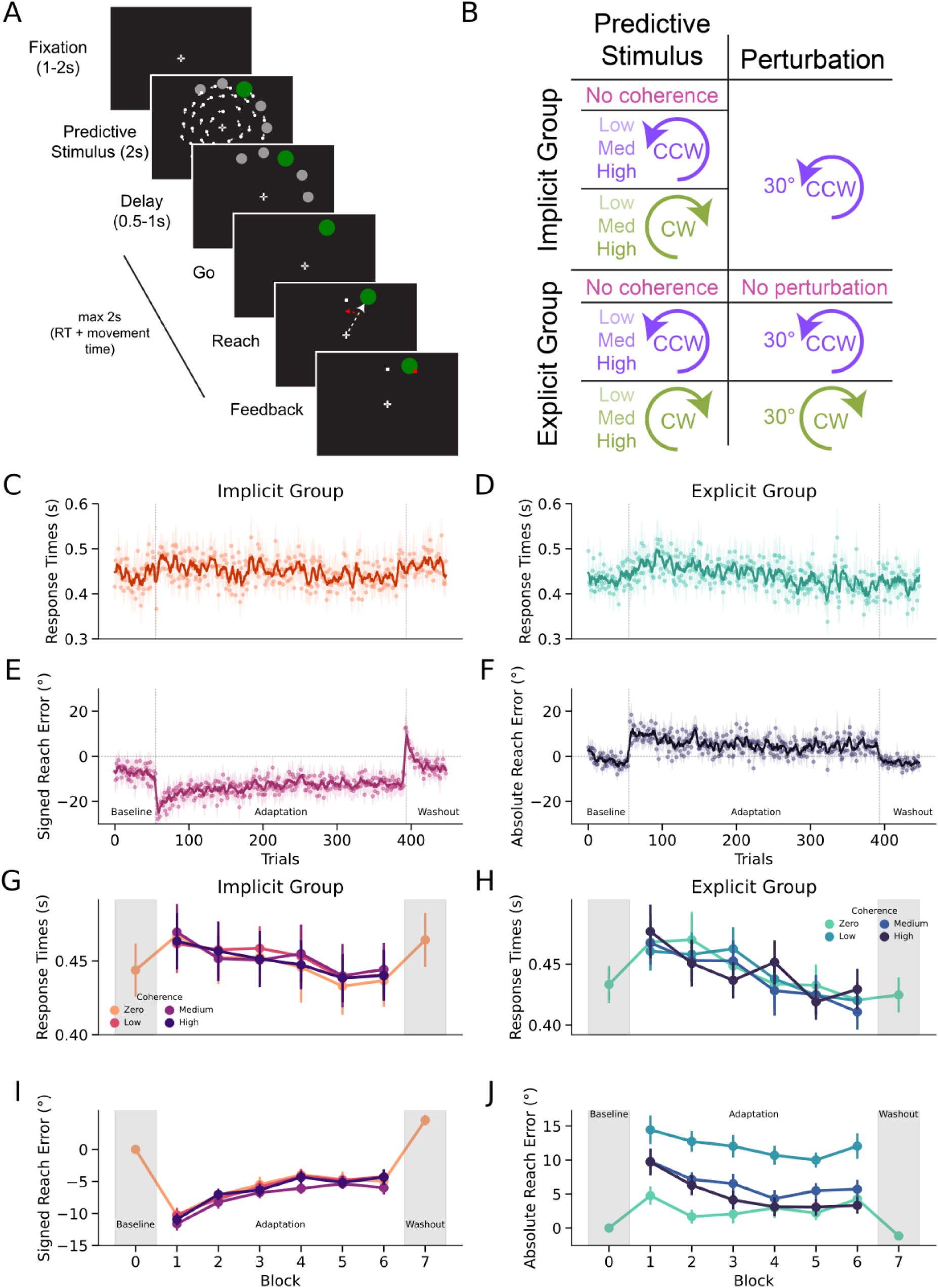
Task design and behavioral dissociation between implicit and explicit learning. **(A)** Trial structure. Each trial began central fixation (1-2 s), followed by presentation of a circular random dot kinematogram (RDK; 2s). Five potential targets were positioned along the RDK edge; one was enlarged and colored green to indicate the reach target. After a brief delay (0.5-1 s), distractor targets disappeared, serving as the go cue. Participants had up to 1 s to move a joystick-controlled cursor to the target, followed by visual feedback. **(B)** Experimental manipulation. RDK motion coherence varied (zero, low, medium, high). Participants were assigned to an implicit or explicit group. During adaptation blocks, the implicit group experienced a constant −30° visuomotor rotation irrespective of RDK motion. In the explicit group, rotation was contingent on RDK coherent motion direction: −30° for counterclockwise (CCW), +30° for clockwise (CW), and 0° for zero coherence. **(C, D)** Response times (RTs) across trials for the implicit and explicit groups. Points indicate group means; shaded areas indicate ± SEM. **(E, F)** Signed and absolute reach error across trials for the implicit and explicit groups (mean ± SEM). **(G, H)** Block-wise response times stratified by coherence level for the implicit and explicit groups (mean ± SEM). **(I, J)** Block-wise signed and absolute reach error, grouped by coherence level for the implicit and explicit groups (mean ± SEM).

### Response time reveals strategic engagement in the explicit group

We first tested whether the task manipulation dissociated implicit from explicit motor learning. Response times (RTs) showed a group × block type interaction (*χ*²(2) = 35.234, *p* < 0.001). In the explicit group, RTs increased during adaptation relative to baseline (*z* = 2.555, *p* = 0.0286) and washout (*z* = 4.615, *p* < 0.001; Figure 1D), consistent with the engagement of a deliberate re-aiming strategy. In contrast, the implicit group showed shorter RTs in the baseline and adaptation blocks relative to washout (baseline: *z* = −4.04, *p* < 0.001; adaptation: *z* = −3.767, *p* < 0.001; Figure 1C). Overall RTs did not differ between groups, indicating that subsequent neural differences cannot be attributed to global task difficulty.

Within adaptation blocks, RTs decreased across blocks in both groups (implicit: *χ²* = 32.2270, *p* < 0.001; explicit: *χ²* = 122.420, *p* < 0.001), consistent with progressive learning. Critically, only the explicit group showed a coherence ⨉ block interaction (*χ*²(3) = 47.430, *p* < 0.001), with slower responses on zero-coherence trials compared to all other coherence levels (all *p* < 0.001), whereas coherence had no effect in the implicit group (Figure 1G-H). Thus, strategic preparation in the explicit group scaled with cue reliability.

### Error dynamics confirm implicit aftereffects and explicit strategy use

Hallmarks of implicit sensorimotor adaptation include initial undercompensation during early adaptation resulting in large reach errors, gradual error reduction across blocks, and a post-adaptation overshoot^18,20^. As the visuomotor rotation was unrelated to the RDK motion for the implicit group, implicit reach errors should be independent of coherence level. Conversely, explicit, top-down re-aiming strategies, driven by RDK cues, should yield coherence level-dependent error patterns. Absolute reach error (i.e., error magnitude) revealed a robust group ⨉ block type interaction (*χ*²(2) = 672.74, *p* < 0.001). In the explicit group, adaptation was associated with increased error relative to baseline (*z* = 18.804; *p* < 0.001) and washout (*z* = 22.171; *p* < 0.001; Figure 1F), reflecting deliberate re-aiming under variable perturbations. By contrast, in the implicit group, error decreased in adaptation relative to baseline (*z* = −3.62; *p* = 0.0009), and increased during washout relative to adaptation (*z* = 4.473; *p* < 0.001), reflecting classic aftereffects of implicit learning^18^.

To directly test for signatures of implicit learning, we examined signed reach error (i.e. the directional error, preserving the sign of the angular deviation relative to the target) in the implicit group. Signed error varied significantly across block types (*χ*²(2) = 922.17, *p* < 0.001), decreasing during adaptation relative to baseline (*z* = −16.406; *p* < 0.001), and reversed sign during washout, producing a significant overshoot relative to both baseline (*z* = 8.577, *p* < 0.001) and adaptation (*z* = 27.622; *p* < 0.001, Figure 1E). Signed error further decreased across successive adaptation blocks (*χ²* = 175.18, *p* < 0.001). This washout overshoot constitutes a canonical aftereffect of implicit sensorimotor recalibration.

### Coherence selective modulates explicit but not implicit adaptation

Within the adaptation blocks, coherence did not influence signed reach error in the implicit group (no main effect or interaction; Figure 1I), consistent with the perturbation being unrelated to the RDK cue. In contrast, the explicit group showed a coherence ⨉ adaptation block interaction (*χ*²(3) = 12.380, *p* = 0.006), with a significant effect of block in low (*z =* −2.962, *p =* 0.0031), medium (*z* = −3.386, *p* < 0.001) and high (*z* = −5.870, *p* < 0.001) coherence trials, and stronger error reduction across blocks at higher coherence levels (high vs. zero: *z* = 3.444, *p* = 0.003; Figure 1J).

These findings demonstrate a clear behavioral dissociation: implicit learners exhibited gradual error reduction and robust aftereffects independent of cue coherence, whereas explicit learners showed strategy-dependent increases in RT and coherence-sensitive error dynamics without aftereffects. This dissociation validates the task manipulation and establishes a behavioral framework for interpreting subsequent neural analyses.

### Beta power and burst rate show distinct sensitivities to learning context

Having established a behavioral dissociation between implicit and explicit learning, we next asked whether beta dynamics during movement preparation and outcome evaluation reflected these differences. We quantified single-trial beta power and detected transient beta burst events from sensors located over sensorimotor cortex, contralateral to the hand used to perform the movement. Analyses focused on two task epochs: a visual epoch time-locked to RDK onset and a motor epoch time-locked to movement completion. All measures were baseline-corrected relative to the pre-stimulus interval.

### Dissociation between beta power and burst rate during adaptation

No group differences were observed during baseline or washout blocks. During adaptation, however, a dissociation emerged in the visual epoch. Prior and around cue onset, the implicit group exhibited stronger beta power suppression than the explicit group (Figure 2A). In contrast, the explicit group showed a selective reduction in overall beta burst rate approximately 1 s after cue onset (Figure 2B). Thus, beta power and burst rate diverged in their sensitivity to learning strategy during movement preparation. In the motor epoch, post-movement beta rebound (PMBR) amplitude did not differ between groups across experimental blocks, indicating that group differences were specific to preparatory dynamics.

**Figure 2.**
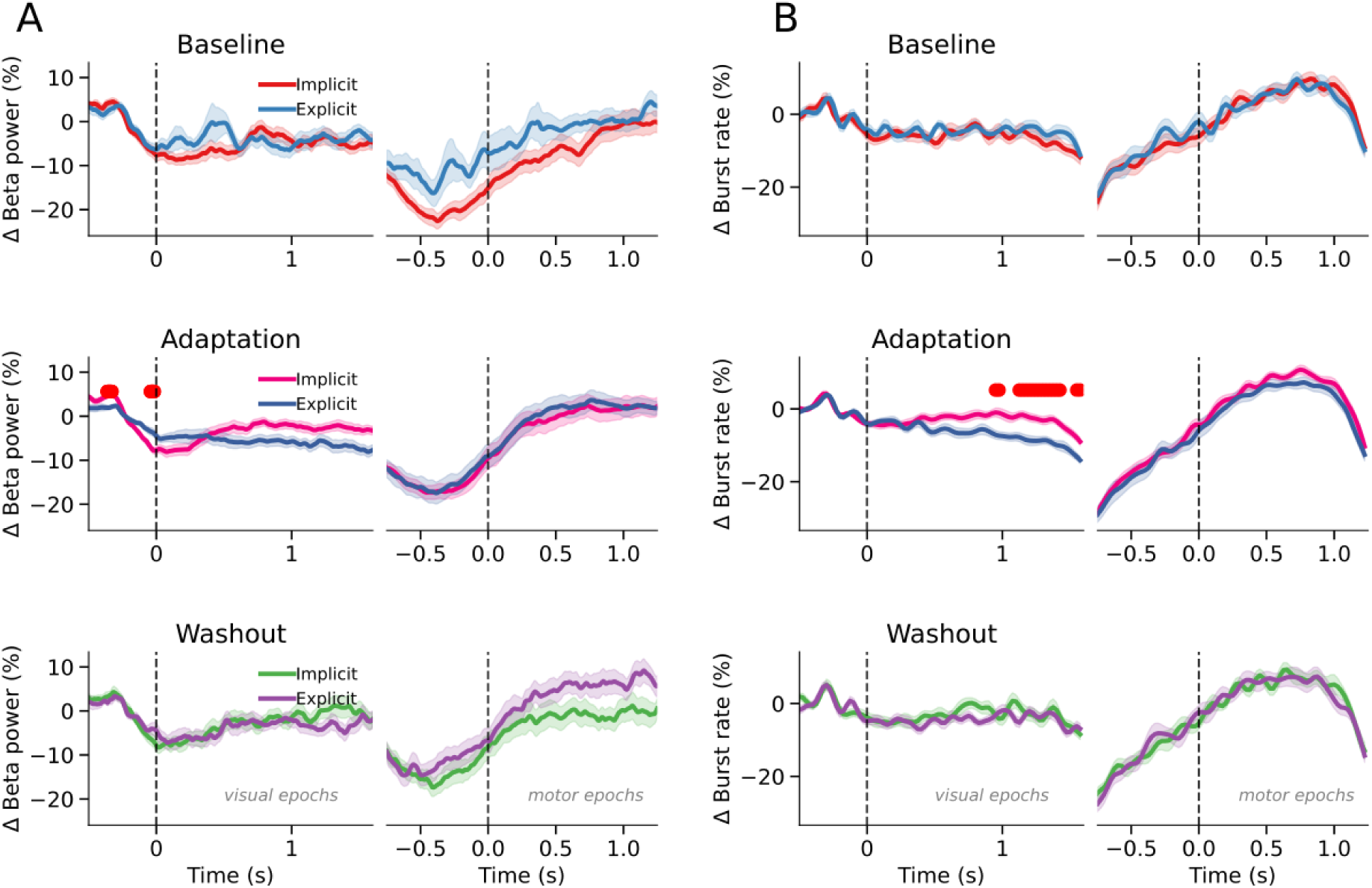
Divergent modulation of beta power and overall burst rate during motor adaptation. **(A)** Baseline-corrected beta-band power (Δ beta power, %) from left-central sensors over contralateral sensorimotor cortex during baseline, adaptation, and washout blocks. Traces are shown separately for implicit and explicit groups, aligned to cue onset (visual epochs) and movement completion (motor epochs; dashed vertical lines at time 0). **(B)** Corresponding baseline-corrected overall beta burst rate (Δ burst rate, %) from the same sensors and task epochs. Red markers indicate time windows with significant between-group differences. Shaded areas denote ± SEM.

### Block-specific modulation of burst rate in the explicit group

When comparing experimental blocks within each group, beta power showed comparable modulation across baseline, adaptation, and washout blocks in both groups (Figure 3A). Burst rate followed a similar pattern in the implicit group. In contrast, the explicit group exhibited a selective reduction in burst rate during adaptation relative to both baseline and washout (Figure 3B), confirming that the adaptation-related decrease in burst rate was both group-specific (Figure 2B) and block type-specific (Figure 3B).

**Figure 3.**
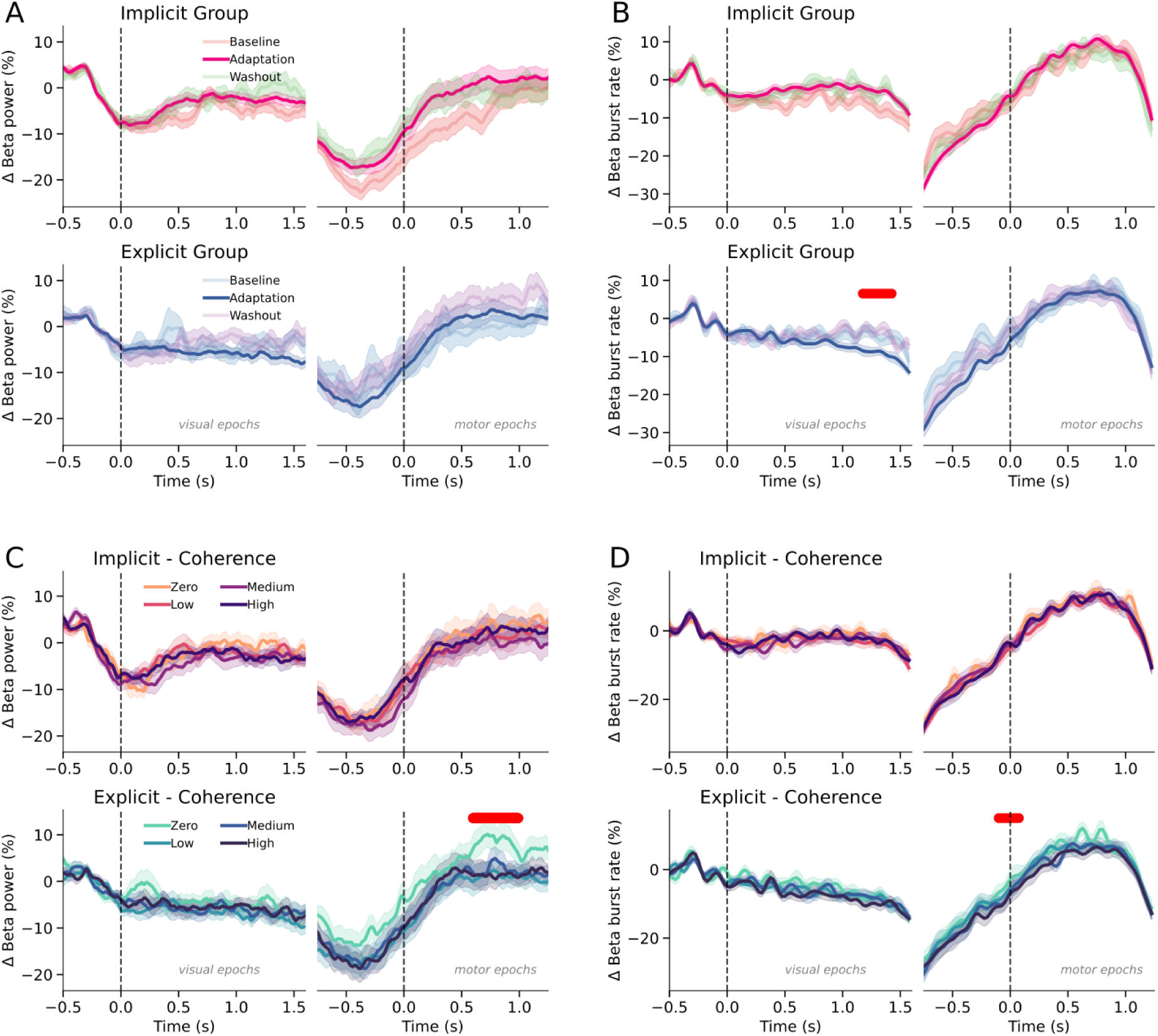
Block- and coherence-dependent modulation of beta power and overall burst rate. **(A)** Baseline-corrected beta power (Δ beta power, %) across baseline, adaptation, and washout blocks, shown separately for the implicit (top) and explicit (bottom) groups. Traces are aligned to cue onset (visual epochs) and movement completion (motor epochs; dashed vertical lines at time 0). **(B)** Corresponding baseline-corrected overall beta burst rate (Δ burst rate, %) across experimental blocks for each group. **(C)** Beta power during adaptation blocks, stratified by motion coherence level (zero, low, medium, high) for the implicit (top) and explicit (bottom) groups. **(D)** Corresponding overall beta burst rate during adaptation blocks, grouped by coherence level. Red markers indicate time windows with significant condition differences. Shaded areas denote ± SEM.

### Coherence sensitivity selectively modulates beta dynamics in the explicit group

We next examined whether beta dynamics tracked the reliability of the predictive cue. In the implicit group, neither beta power nor burst rate varied with coherence level during adaptation, consistent with the cue being behaviorally irrelevant. In the explicit group, however, beta dynamics were coherence-sensitive. High-coherence trials were associated with reduced burst rate during movement preparation (Figure 3D), whereas zero-coherence trials elicited greater beta power and burst rate during the post-movement rebound (Figure 3C and 3D).

These findings reveal a functional dissociation: beta power primarily indexed preparatory suppression in the implicit group, whereas burst rate selectivity tracked strategic engagement and cue reliability in the explicit group. This divergence across block and learning context indicates that beta power and burst rate are not interchangeable metrics and suggests that averaging across bursts may obscure distinct neural processes underlying motor adaptation.

### Waveform-resolved burst dynamics reveal functional heterogeneity

We next examined whether beta burst morphology differed between implicit and explicit learning contexts, first using conventional time-frequency metrics and then assessing waveform variability in the time domain.

### Small but consistent differences in time-frequency features

We compared standard burst features between groups using linear mixed-effects models (LMMs) (group as a fixed effect; subject-specific random offsets). Reliable but small group differences were observed across both visual and motor epochs. Burst duration differed between groups (visual: χ²(2) = 93.93, *p* < 0.001; motor: χ²(2) = 437.94, *p* < 0.001; Figure 4A, 4C), with bursts in the explicit group lasting ∼ 2 ms longer on average. Peak amplitude also differed (visual: χ²(2) = 590.29, *p* < 0.001; motor: χ²(2) = 4210, *p* < 0.001; Figure 4B, 4D), as did frequency span (visual: χ²(2) = 55.02, *p* < 0.001; motor: χ²(2) = 265.36, *p* < 0.001; Figure 4E, 4G), although the absolute differences in frequency span were minimal (∼ 0.03 Hz). Peak frequency did not differ between groups in either epoch (visual: χ²(2) = 1.01, *p* = 0.60; motor: χ²(2) = 2.17, *p* = 0.34; Figure 4F, 4H).

**Figure 4.**
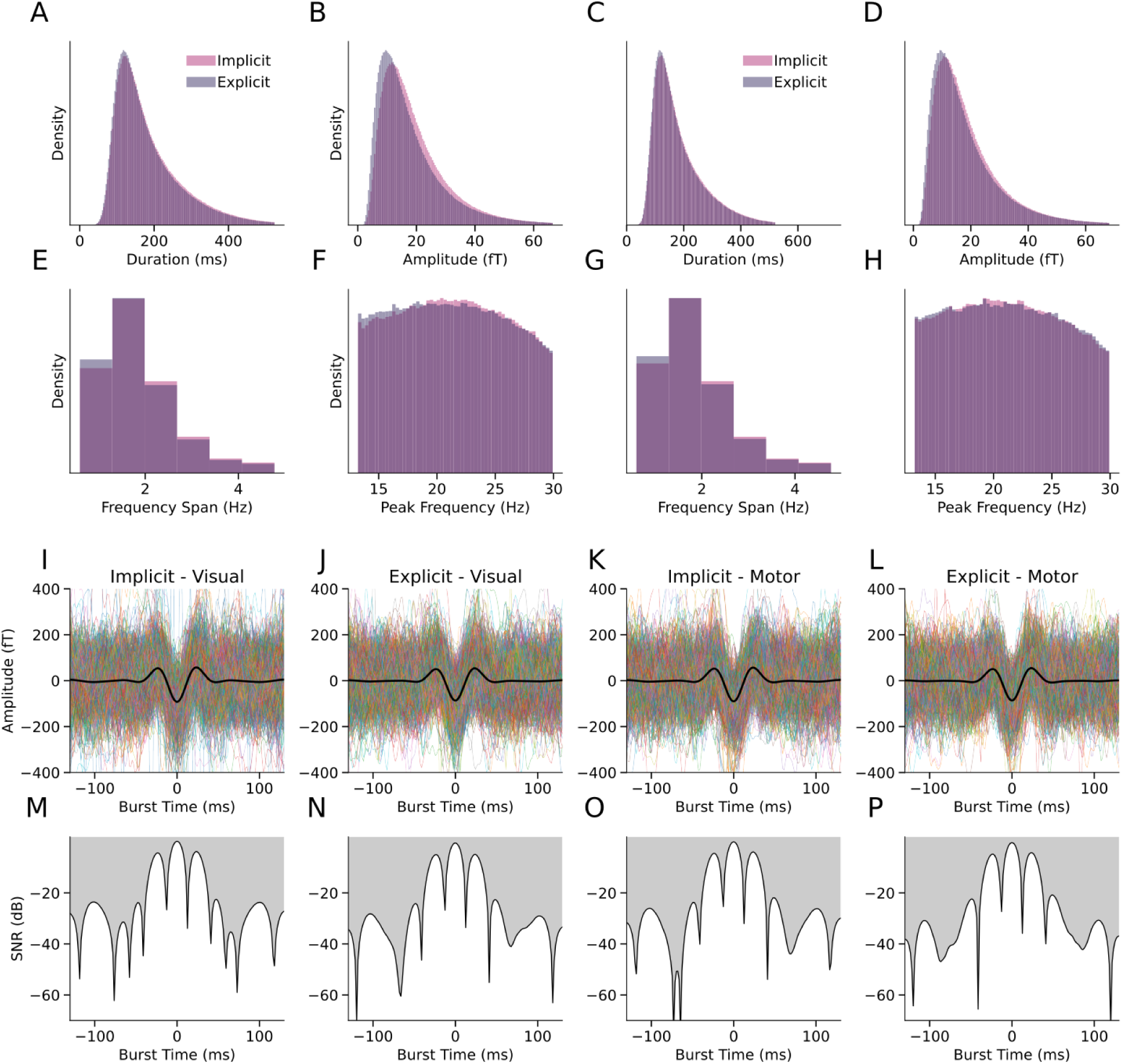
Time-frequency features and waveform structure of beta bursts. **(A-D)** Distributions of burst duration and peak amplitude for implicit and explicit groups in visual and motor epochs. **(E-H)** Corresponding distributions of frequency span and peak frequency across epochs. **(I-L)** Burst waveforms for the implicit and explicit groups during visual and motor epochs. Thin colored lines represent individual bursts; the thick black trace indicates the mean waveform across bursts. **(M-P)** Signal-to-noise ratio (SNR, dB) time courses for bursts in the same visual and motor epochs, quantifying burst-to-burst variability around the mean waveform.

Despite these small time-frequency differences, the mean burst waveform and its signal-to-noise ratio were indistinguishable between groups or epochs (Figure 4I-P), indicating that learning context did not alter the stereotyped burst waveform shape. We therefore asked whether variability within the waveform space, rather than the mean waveform, was differentially modulated across task conditions.

### Principal components define task-modulated waveform dimensions

Because beta bursts are brief, nonstationary events whose waveform shape may vary even when spectral characteristics are similar ^8,21–24^, we analyzed burst waveforms directly in the time domain using principal component analysis (PCA; Figure 5A). A permutation procedure retained only components explaining variance beyond noise fluctuations. Four components (PC7-PC10) showed systematic modulation across task epochs, indicating that specific waveform features changed in prevalence over time (Figure 5C).

**Figure 5.**
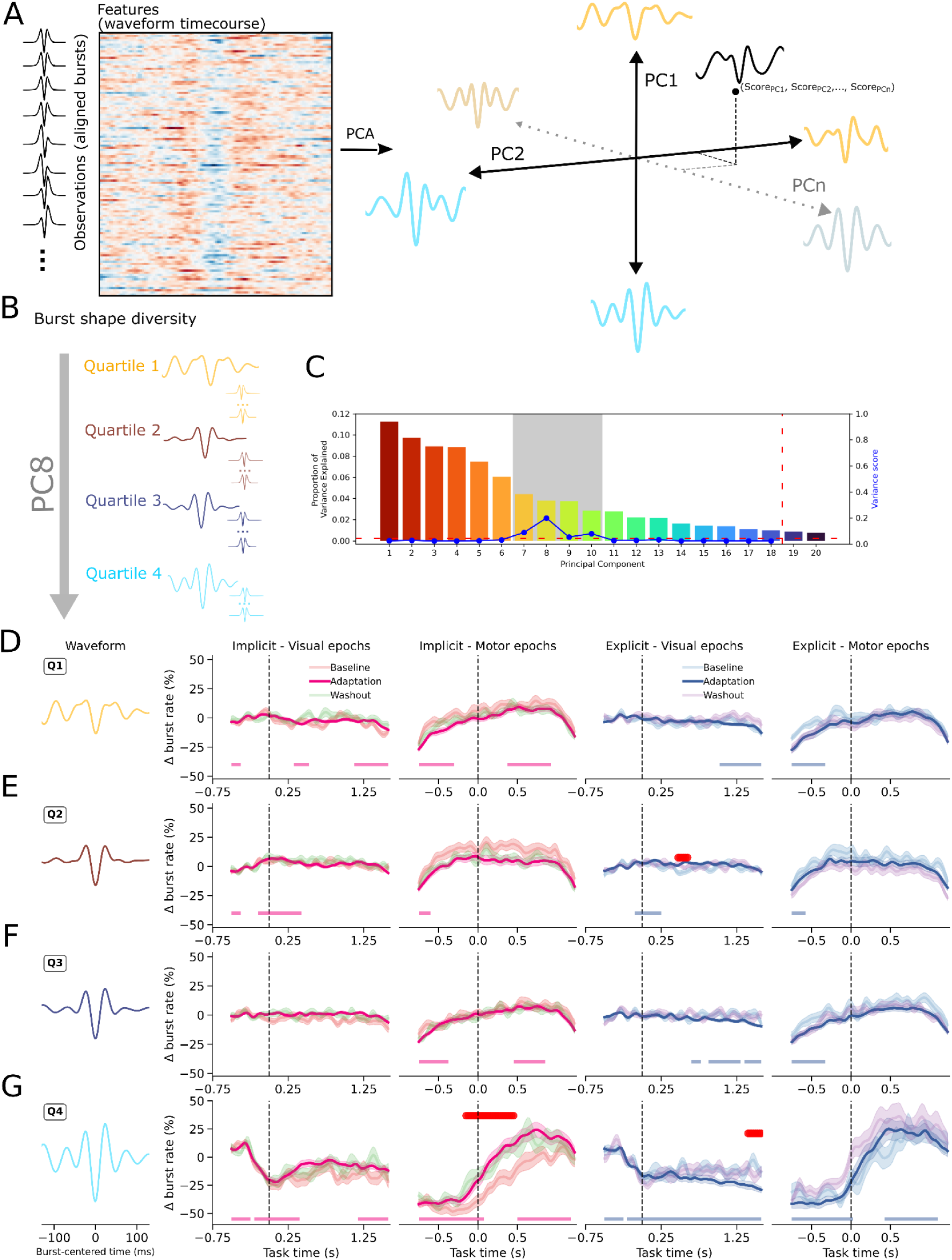
Principal component decomposition reveals waveform-specific burst dynamics. **(A)** Schematic of the PCA-based approach. Individual burst waveforms (aligned to burst peak) were arranged as observations × time points and decomposed into orthogonal principal components (PCs), defining continuous dimensions of waveform variability. **(B)** Burst shape diversity was quantified by projecting each burst onto a given PC axis and dividing the distribution into quartiles (Q1-Q4), thereby defining burst types occupying different positions along that waveform dimension (illustrated for PC8). **(C)** Variance explained by each principal component. The dashed red vertical line marks the cutoff determined by permutation testing; components to the left of this threshold were retained for further analyses. The blue line indicates the variance score, defined as the difference in variance of the mean component score time course between visual and motor epochs. The shaded region highlights components exceeding the significance threshold and showing task-related modulation. **(D-G)** Baseline-corrected burst rates (Δ burst rate, %) separated by PC8 quartile (Q1-Q4), shown across visual and motor epochs, groups (implicit, explicit), and experimental blocks (baseline, adaptation, washout). Bottom markers indicate time periods where burst rates during adaptation significantly differed from zero. Red markers indicate significant condition differences. Shaded areas represent ± SEM.

Each burst was assigned a score along each component axis (Figure 5B). To characterize burst rate dynamics, we divided the score distribution for each component into quartiles, defining *burst types* corresponding to different positions along a continuous waveform dimension (Figure 5B). Primary analyses focused on PC8, which exhibited the strongest task-related modulation (Figure 5C; results for PC7, PC9 and PC10 are provided in the Supplementary Materials, Figure S1-3).

### Distinct burst types exhibit dissociable rate profiles

Bursts divided along PC8 quartiles displayed significantly different rate dynamics. One-sample cluster tests revealed distinct modulation patterns during visual epochs of adaptation blocks. Q2 bursts showed post-stimulus rate increases most prominent in the implicit group (Figure 5E), whereas Q4 bursts exhibited pronounced suppression below baseline following cue onset, also particularly in the implicit group (Figure 5G). Q1 bursts showed a later reduction in both groups (Figure 5D), and Q3 bursts showed no sustained modulation in the implicit group but a late reduction in the explicit group (Figure 5F). Block-wise comparisons revealed condition-specific modulation primarily in Q2 and Q4 in the explicit group, with lower burst rates during baseline relative to adaptation and washout blocks in Q2 (Figure 5E), and lower burst rates during adaptation trials in Q4 (Figure 5G). Crucially, direct comparison revealed significantly lower Q3 and Q4 burst rates in the explicit compared to the implicit group during adaptation trials (Figure 6C,D).

**Figure 6.**
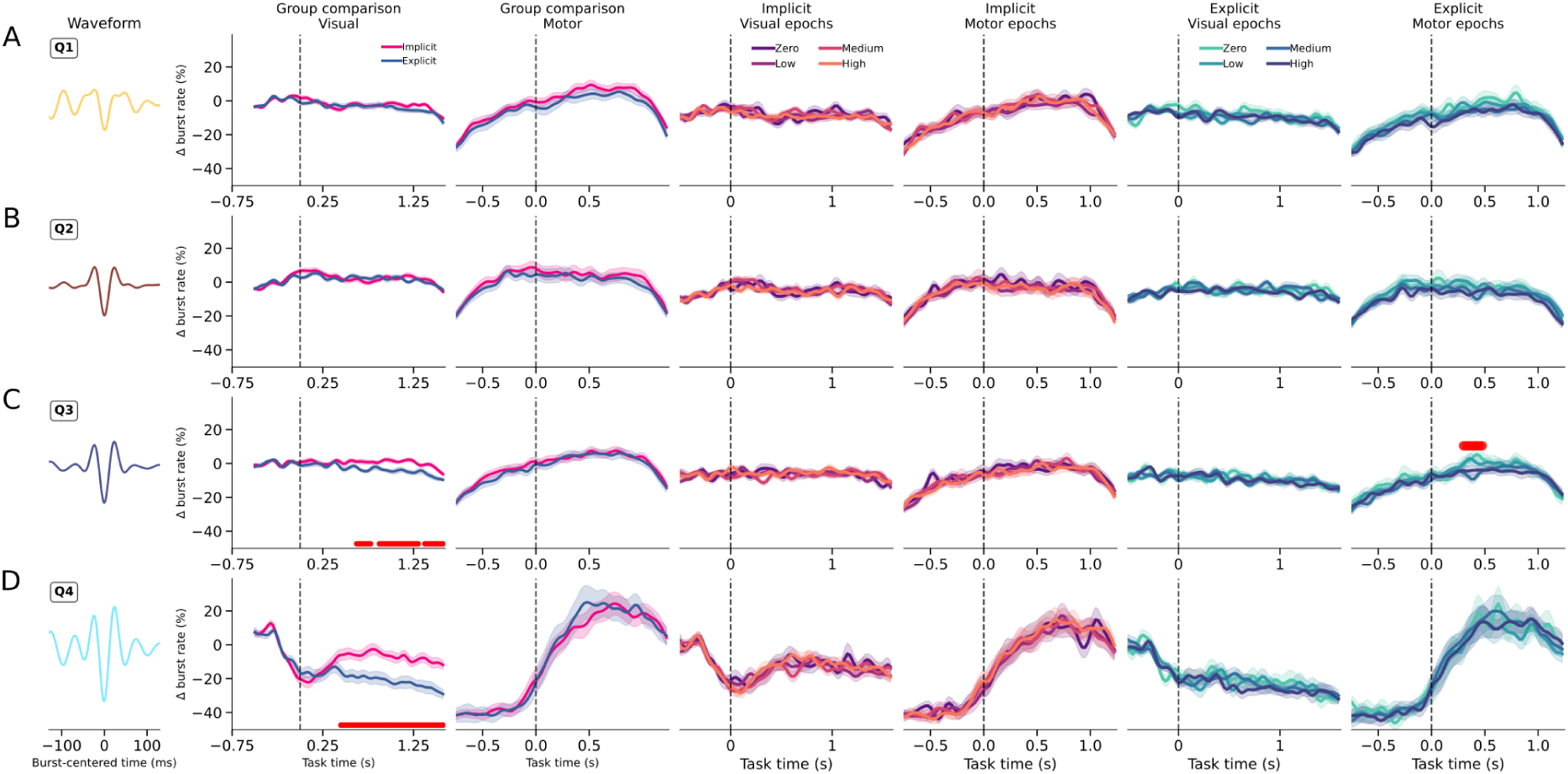
Coherence-dependent modulation of waveform-defined burst types during adaptation. **(A-D)** Baseline-corrected burst rates (Δ burst rate, %) within adaptation blocks, stratified by motion coherence level (zero, low, medium, high) and shown separately for implicit and explicit groups. Panels correspond to PC8 waveform quartiles (Q1-Q4), with traces aligned to cue onset (visual epochs) and movement completion (motor epochs; dashed vertical lines at time 0). Red markers denote time windows with significant coherence-related differences. Shaded areas represent ± SEM.

In motor epochs, two time windows are of interest: during movement and post-movement. Overall, all burst types showed a rate reduction during movement in both groups, with no differences across block type. Following movement completion, Q1, Q3, and Q4 burst rates increased gradually in the implicit group (Figures 5D, F, G), whereas only Q4 bursts showed a post-movement increase in the explicit group (Figure 5G). Block-wise comparisons further revealed that Q4 bursts exhibited the strongest contextual modulation: in the implicit group, adaptation and washout trials were associated with higher peri-movement burst rates relative to baseline (Figure 5G).

### Coherence-dependent modulation of waveform-defined burst types

During adaptation, coherence-level analyses revealed waveform-specific effects consistent with earlier power and overall burst rate findings. In the explicit group, zero-coherence trials elicited higher post-movement burst rates in Q3 trials (Figure 6C). No coherence-related differences were observed in the implicit group.

These results indicate that although the average beta burst waveform is preserved across learning contexts, variability along specific waveform dimensions is differentially modulated by task epoch, learning block, and cue reliability. Beta modulation during adaptation therefore operates within a structured, low-dimensional space of burst morphologies rather than through uniform changes in overall burst rate.

### Waveform-specific burst dynamics predict trial-by-trial error

To link neural dynamics to behavior, we quantified the relationship between beta activity (power, overall beta burst rate, and waveform-specific burst rates) and trial-wise performance (response time and error magnitude). Linear mixed-effects models were fitted with behavior as the dependent variable, beta metric and group as fixed effects, and subject-specific random intercepts. A sliding-window approach tracked the temporal evolution of these effects, extracting the beta coefficient and associated *p*-value at each time point. Positive coefficients indicate that larger behavioral values were associated with higher beta activity; negative coefficients indicate an inverse relationship.

Neither beta power nor overall burst rate predicted response time in either epoch (Figure 7A-B). In contrast, error magnitude showed a robust relationship with post-movement beta power in both groups. From 0.175 - 1.075 s after movement completion, larger errors were associated with a reduced beta rebound (all *p* < 0.001, Figure 7A). No corresponding relationship was observed for overall beta burst rate (Figure 7B), indicating that power-error coupling was not captured by aggregate bust counts.

**Figure 7.**
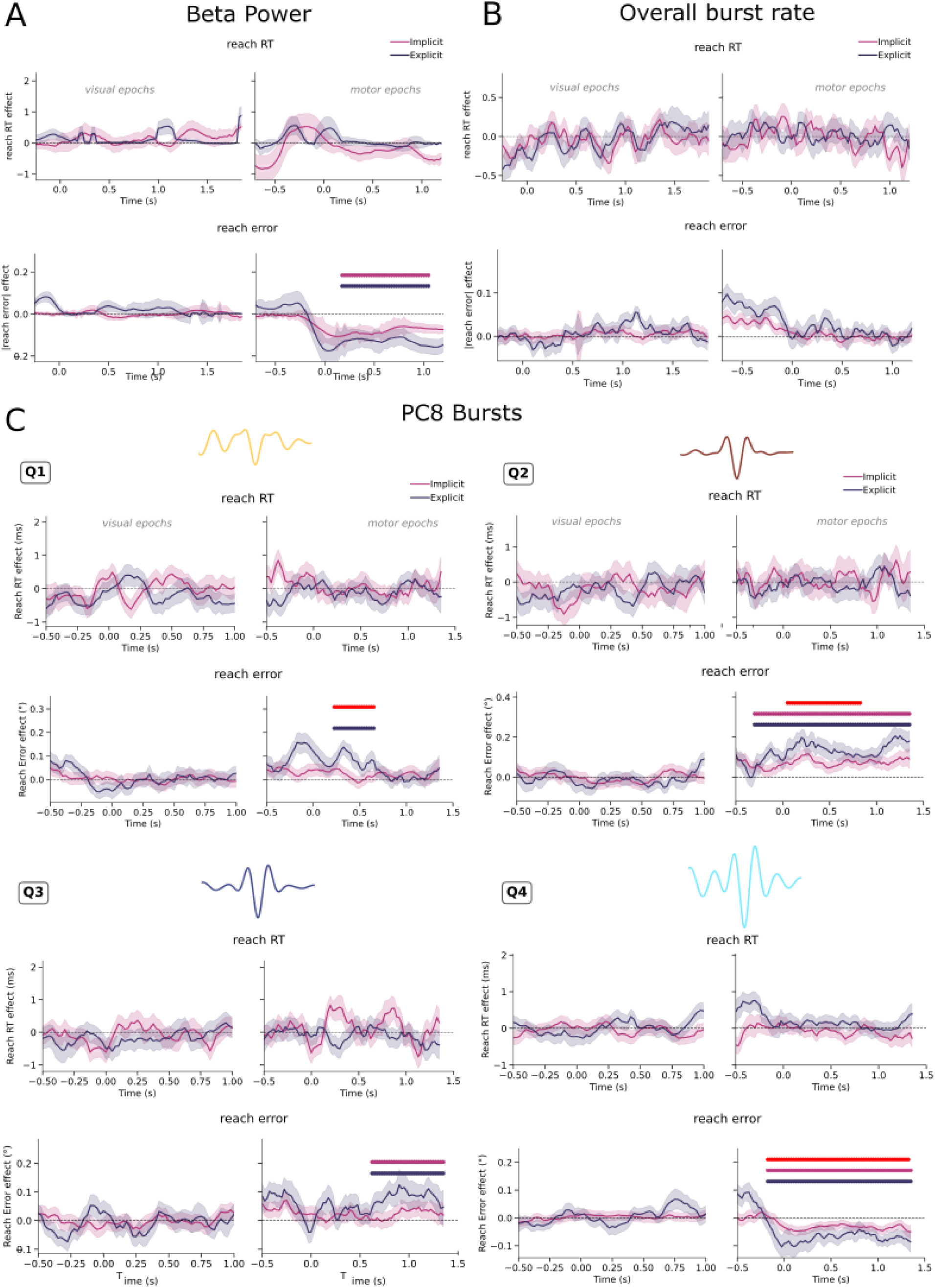
Waveform-specific beta dynamics predict trial-by-trial behavior. Sliding-window linear mixed-effects analyses relating neural activity to behavioral performance. For each time window, the standardized effect size (β coefficient) of the neural predictor is plotted over time, separately for visual and motor epochs. Positive values indicate a positive relationship between the beta metric and behavior; negative values indicate an inverse relationship. **(A)** Relationship between beta power and reach response time (RT) and reach error. **(B)** Relationship between overall burst rate and reach RT and reach error. **(C)** Relationship between waveform-sorted burst rate along PC8 (quartiles Q1-Q4) and RT and reach error. Purple markers denote significant effects in the implicit group; blue markers denote significant effects in the explicit group; red markers indicate significant group × neural predictor interactions. Shaded areas represent ± SEM.

Waveform-resolved analyses revealed a structured dissociation across burst types. Q4 bursts reproduced the beta-power pattern, showing a negative relationship between error and burst rate during motor epochs (*χ*²(1) = 47.74, *p* < 0.001; −0.17-1.35 s). Both groups exhibited significant negative slopes (explicit: *z* = −5.85, *p* < 0.001; implicit: *z* = −3.50, *p* < 0.001), with a stronger effect in the explicit group (interaction: *χ*²(1) = 6.44, *p* = 0.011; Figure 7C).

In contrast, Q1 and Q2 bursts showed error-rate relationships. Q1 bursts exhibited a group difference (*χ*²(1) = 7.62, *p* = 0.006; 0.23-0.65s; Figure 7C), with a significant positive slope in the explicit group (*z* = 4.18, *p* < 0.001) but not in the implicit group (*z* = 0.84, *p* = 0.40). Q2 bursts showed a robust positive relationship with error (*χ*²(1) = 56.49, *p* < 0.001; −0.30-1.35 s) in both groups (explicit: *z* = 4.70, *p* < 0.001; implicit: *z* = 4.48, *p* < 0.001), again stronger in the explicit group (*χ*²(1) = 4.15, *p* = 0.042). Q3 bursts exhibited a later positive relationship (*χ*²(1) = 9.14, *p* = 0.003; 0.62-1.35 s) without group differences.

Thus, while aggregate beta measures obscure behaviorally relevant structure, waveform-defined burst types show opposing and temporally specific relationships with error magnitude. These results indicate that beta-behavior coupling resides within distinct burst motifs rather than in overall burst rate or power, supporting a functionally heterogeneous view of beta dynamics.

## Discussion

We asked whether variability in beta burst waveforms reflects functionally distinct neural processes during motor learning. By distinguishing bursts based on their waveforms, we show that specific burst types exhibit opposing and temporally specific relationships with behavioral error. In addition, particular waveform-defined burst populations show context-dependent modulation during movement preparation, differentiating implicit and explicit learning processes. In contrast, conventional measures of beta activity, such as overall power and burst rate, showed reduced specificity of these dynamics, reinforcing our position that beta bursts should not be treated as a homogeneous phenomenon ^22^.

### Behavioral differences between implicit and explicit learning

The task manipulation successfully dissociated implicit from explicit learning. The implicit group showed classic signatures of sensorimotor adaptation: gradual reach error reduction during adaptation followed by robust washout aftereffects, with performance independent of dot coherence in the RDK. In contrast, the explicit group exhibited increased response times during adaptation and coherence-dependent error patterns without aftereffects, confirming strategic re-aiming based on the predictive cue (Figure 1). This behavioral dissociation enables us to interpret the neural results in terms of learning context and assess whether brain responses differ between implicit and explicit strategies.

### Pre-movement beta dynamics

Previous work has shown that the pre-movement beta decrease is closely linked to motor planning demands, but its precise functional role, whether facilitating motor readiness ^25,26^, enhancing sensorimotor integration ^27,28^, or reflecting attentional engagement, remains debated ^29,30^. Our results speak to this question: implicit learning was associated with stronger sensorimotor beta desynchronization around RDK onset (Figure 2A). Because the RDK was uninformative in the implicit condition, this enhanced desynchronization might reflect increased reliance on sensory feedback, as the motor system continuously integrates prediction errors to drive forward model updating ^31–33^. In contrast, explicit learning was characterized by a reduced beta burst rate during the later preparation phase (∼1 s after RDK onset; Figure 2B, 3B), potentially reflecting the selection of the appropriate motor command consistent with the re-aiming strategy, a process attributed to inverse model computations, where the desired movement outcome is mapped onto the required motor output ^34^. This temporal and metric-specific dissociation indicates that beta power and burst rate are not interchangeable metrics, each capturing distinct aspects of preparatory dynamics during adaptation.

Waveform-resolved burst analyses clarify this preparatory dissociation. Following RDK onset, distinct burst populations exhibited opposing dynamics: Q2 bursts increased in rate, Q4 bursts showed an early suppression, and Q1 and Q3 bursts showed later reductions (Figure 5). Crucially, Q3 and Q4 burst rates were selectively reduced in the explicit group during the visual epoch (Figure 6C,D), indicating that specific waveform-defined motifs are differentially engaged when participants rely on strategic re-aiming. These effects suggest that preparatory beta dynamics are not simply globally suppressed or enhanced, but are reweighted within a structured burst repertoire depending on learning strategy.

### Movement-related beta

Our epoching around reach completion (i.e., motor epochs) allowed us to dissociate beta dynamics linked to movement execution (pre-0) from those related to post-movement evaluation (post-0). Across groups, beta power, overall burst rate and waveform-defined burst types showed a robust suppression prior to movement completion, consistent with the well-established movement-related beta decrease reflecting motor execution ^1,28,35^. This movement-related reduction did not differ across learning context or block type, suggesting that core execution-related beta dynamics are preserved across learning contexts. No reliable relationship was observed between pre-movement completion beta activity and trial-by-trial behavior (Figure 7), indicating that execution-related beta suppression was not predictive of performance variability.

### The two faces of the beta rebound

In contrast to preparatory and movement-related beta reduction, the beta rebound has been put forward as either an active process to maintain the *status quo* of motor planning ^2^ or a comparison between the planned and actually performed movement, reflecting the certainty of feedforward action signals based on internal models ^36^. For example, previous studies have shown that PMBR amplitude is positively modulated by the confidence in the correctness of our actions ^36^, and is reduced when large errors are committed ^37^. Conversely, others have suggested that the amplitude of the beta rebound is linked to the magnitude of errors, showing smaller beta power for smaller errors, hence potentially encoding increased response inhibition ^38^. These two theories make different predictions regarding the relationship between beta activity and errors: the confidence-based account predicts a decrease in beta power as errors increase, while the response inhibition account suggests that beta power should instead rise with greater errors.

Our findings provide support for the confidence-based account of the post-movement beta rebound. Across groups, larger reach errors were associated with a reduced post-movement beta power, emerging approximately 175 ms after movement completion and extending beyond 1 s (Figure 7A). As performance worsened, the amplitude of the rebound decreased, consistent with the idea that PMBR reflects the certainty or stability of internal forward model predictions.

Crucially, this error-beta relationship was not captured by overall burst rate (Figure 7B). Instead, waveform-resolved analyses revealed that the negative relationship between beta activity and error was primarily driven by a specific burst population (Q4). Q4 burst rate closely mirrored the beta power effect, showing a negative association with error magnitude in both groups, stronger in the explicit condition. In contrast, other burst types (Q1, Q2, Q3) showed positive relationships with error, indicating that distinct waveform-defined motifs carry opposing behavioral signatures within the same post-movement window. Our waveform-resolved analyses therefore suggest that apparently conflicting accounts of the beta rebound may reflect the aggregation of multiple burst motifs with opposing functional signatures. When considered separately, Q4 bursts align with confidence-based interpretations of PMBR, whereas Q1-Q3 bursts exhibit dynamics consistent with ongoing corrective or evaluative processing.

Together, these findings indicate that the post-movement beta rebound is not a unitary signal but an emergent composite of multiple burst populations with distinct and sometimes opposing computational roles.

## Methods

### Participants

The experiment involved 38 healthy, right-handed participants (25 females, aged 20-35 years, M = 26.69, SD = 4.11 years) with normal or corrected-to-normal vision. The study followed the guidelines of the Declaration of Helsinki, and all participants provided written consent approved by the regional ethics committee for human research (CPP Est IV - 2019-A01604-53).

### Behavioral Task

Participants were asked to perform ballistic joystick-based movements with their right hand to reach a visual target. At the beginning of each trial, participants were instructed to fixate on a small (0.6° ⨉ 0.6°) central target comprising both a bullseye and crosshairs (see Figure 1;^39^). After a variable delay ranging from 1 to 2 seconds, a circular random dot kinematogram (RDK) was displayed, exhibiting coherent motion in either a clockwise or counter-clockwise direction. Five circular targets were positioned along the outer edge of the RDK, spaced at 30° intervals from −120° to 0°, relative to the fixation point. In each trial, one of the targets located at either −90°, −60°, or −30° was larger (3.25°) and colored green, indicating that participants were to reach for that specific target upon receiving the go cue. The remaining targets were smaller (1.625°) and gray in color (Figure C). After 2 seconds, the RDK disappeared, leaving only the gray potential reach targets and the green instructed target visible. Following a variable delay period ranging from 0.5 to 1 second, the gray targets disappeared, leaving only the green instructed target visible. This served as the go cue, signaling the participant to make a ballistic reach for this target using the joystick. The joystick controlled the position of a small (0.5 ⨉ 0.5°) square white cursor. Trials ended under three conditions: if the distance between the cursor and the fixation target surpassed that of the center of the instructed target (set at 7°), if the movement was initiated ahead of the go cue, or if the reach was not performed fast enough (i.e., within 1 second after the go cue). Feedback was then given on the final position of the cursor as a red dot, which was on screen for 2 s. Following each trial, there was a random inter-trial interval lasting between 1.5 and 2 seconds, during which participants were instructed to return the cursor to the fixation target.

Participants were divided into two groups (implicit and explicit). In each trial, the RDK exhibited different degrees of clockwise or counterclockwise coherent motion, or no coherent motion at all. The visual position of the cursor was perturbed by rotating it by −30°, 0°, or 30° relative to the hand position, based on the participant group and trial condition. For the implicit group (*N* = 18), the visual position of the cursor was rotated by −30° in every trial. Thus, there was no relationship between the direction of motion coherence in the RDK and the visuomotor rotation. However, participants in this group could implicitly adapt to the constant rotation. For the explicit group (*N* = 20), the visuomotor rotation was in the direction of coherent motion of the RDK (−30° for counterclockwise coherent motion, 30° for clockwise motion, and no rotation for no coherent motion). Consequently, participants could predict the direction of the perturbation from the RDK and compensate for it with varying levels of difficulty.

The RDK was displayed within a circular window spanning 7° centered on the fixation point, comprising 200 white dots, each with a diameter of 0.3° and moving at a speed of 10°/s. In each trial, a specific percentage of the dots (determined by the motion coherence level) moved coherently around the aperture’s center in either a clockwise or counterclockwise direction, at a distance ranging from 2° to 7° from the fixation target. The remaining dots followed random directions but maintained a consistent trajectory per dot. The lifetime of each dot followed a normal distribution (*M* = 416.67 ms, *SD* = 83.33 ms).

Motion coherence levels were individually calibrated for each participant using an adaptive staircase procedure (QUEST; Watson and Pelli, 1983) to identify the coherence level at which they achieved 82% accuracy across a block of 40 trials at the start of each session. During this block, participants were required to indicate clockwise or counterclockwise motion coherence by pressing the left or right button of a button box, respectively. The coherence level obtained from this procedure was categorized as low, while 150% and 200% of it were considered medium and high, respectively.

Participants underwent 1-3 brief training blocks consisting of 12 trials each. During these sessions, the RDK exhibited no coherent motion, and there was no visuomotor perturbation. Throughout these blocks, the position of the joystick-controlled cursor was visible during the whole trial (online feedback). When the reach distance exceeded the center of the instructed target (7°), the cursor turned red. Additionally, a brief message was displayed for 2 seconds, indicating if participants initiated the reach too early, failed to complete it quickly enough, or deviated too far (> 1.625°) from the target’s center. Once participants achieved at least 75% accuracy within a training block, they progressed to a single block of 56 trials. Similar to the training blocks, this block had no coherent RDK motion or visuomotor rotation. However, the cursor disappeared if its distance from the fixation target exceeded 1° and reappeared, turning red upon reaching the extent of the target (endpoint feedback). After each trial, a brief feedback message was displayed for 2 seconds regarding the timing and accuracy of the reach.

Subsequently, both participant groups completed seven blocks of 56 trials each, with visuomotor rotational perturbation. In these trials, the RDK was presented with varying levels of coherent motion (zero, low, medium, or high) in either clockwise or counterclockwise directions. For the explicit group, the direction of the subsequent visuomotor perturbation (0, −30, or 30°) corresponded to the direction of coherent motion in the RDK. Conversely, for the implicit group, the visuomotor rotation was −30° in every trial, regardless of the RDK’s coherent motion direction. Finally, both groups completed one washout block consisting of 56 trials without coherent motion or visuomotor rotation. The task was implemented using Psychtoolbox (v3.0.16;^40,41^) and executed using Matlab R2017b.

### Behavioral metrics

#### Response Times

We extracted the response times as the time between the appearance of the go cue and the start of the ballistic movement toward the target.

#### Error Measures

To assess participants’ ability to adapt to perturbations, we quantified **r**each error, defined as the angular difference between the final cursor position and the target direction. We first computed absolute reach error (i.e., error magnitude), defined as the absolute difference between the true target angle and the actual reaching angle, while accounting for the applied perturbation. Absolute reach error provides a unified metric of performance accuracy that allows direct comparisons between the implicit and explicit groups across block types. In the implicit group, where the perturbation was constant throughout adaptation (30° irrespective of the visual stimulus), error magnitude reflects deviations from the expected correction. In the explicit group, where perturbation direction varied across trials as a function of the RDK, error magnitude similarly captures deviations in reaching accuracy. To directly test for signatures of implicit learning, we additionally analyzed signed reach error (directional error) in the implicit group only. Because perturbation direction was fixed in this group, directional changes in reach error can be meaningfully interpreted as adaptation and post-adaptation aftereffects.

### MEG

Before coming to the MEG session, each participant underwent an MRI session using a 3 T Siemens Prisma system (Erlangen, Germany). A T1-weighted scan was obtained using a magnetization-prepared rapid gradient-echo (MPRAGE) pulse sequence with a 1 mm isotropic voxel size (256 ⨉ 256 ⨉ 256 voxels). The scan parameters were as follows: repetition time (TR) of 2100 ms, echo time (TE) of 3.33 ms, inversion time (TI) of 900 ms, and a GRAPPA factor of 3. For co-registration of the MRI and MEG data, vitamin E tablets were placed at the nasion and the left and right ear canals.

We utilized the 1 mm T1 MRI volumes to construct individualized foam head-casts for each participant to reduce between-session co-registration error and within-session head movement (i.e., high-precision MEG^42,43^). The scalp surface was extracted from the T1 volume using FreeSurfer (v6.0.0;^44^) and used as a mold for the inner surface of the head-cast. The outer surface was defined by a 3D model of the MEG dewar. These surface models were then aligned relative to a 3D model of the MEG dewar using Rhinoceros 3D (https://www.rhino3d.com) to minimize the distance between the scalp and the sensors without obstructing the participant’s view. The final model was printed using a Raise 3D N2 Plus 3D printer (https://www.raise3d.com). The 3D printed model was placed inside a replica of the MEG dewar, and the space between the head model and the dewar replica was filled with polyurethane foam (Flex Foam-it! 25; https://www.smooth-on.com) to create the participant-specific head-cast, into which the fiducial coils were placed during scanning.

MEG data were recorded using a 275-channel Canadian Thin Films (CTF) MEG system equipped with superconducting quantum interference device (SQUID)-based axial gradiometers (CTF MEG Neuro Innovations, Inc., Coquitlam, Canada) in a magnetically shielded room. Participants were positioned supine during the recording. Visual stimuli were presented on a screen (∼80 cm from the participant) using a video projector (Propixx VPixx, VPixx Technologies Inc., Canada) with a refresh rate of 120 Hz, and participants responded using a joystick (NATA Technologies, Canada). The MEG data were continuously digitized at a sampling rate of 1200 Hz. Eye movement data were captured using an Eyelink 2000 eye tracker (SR Research, Ontario, Canada), which tracked monocular eye movements at 1000 Hz and was calibrated before recording began.

Data preprocessing was primarily performed using the MNE-Python toolbox (v0.23.4^45^; Python v3.9.7). MEG data were downsampled to 600 Hz and filtered with a low-pass 120 Hz zero-phase FIR filter with a Hamming window. Line noise (50 Hz) was removed using an iterative version of the Zapline algorithm^46^ implemented in the MEEGKit package (https://nbara.github.io/python-meegkit/ v0.1.2), with a 20 Hz window for polynomial fitting and a 5 Hz window for noise peak removal and interpolation. Independent Component Analysis (ICA, InfoMax, 25 components) was performed on a band-pass filtered (1 to 60 Hz) copy of the MEG data using the scikit-learn library (v0.24.2;^47^) to isolate ocular movement and cardiac-related artifacts. Eye-tracking data were cropped and resampled to match the MEG signal, and ocular artifacts were identified by correlating each of the 25 components with horizontal and vertical eye movement signals. Blink detection was achieved by thresholding vertical eye movement beyond the vertical resolution of the screen. Components were classified as ocular movement artifacts if all four correlation coefficients (before and after blink removal) exceeded r = 0.15, with an average correlation above r = 0.25. Cardiac artifacts were identified by applying the ECG R peak detector (https://github.com/berndporr/py-ecg-detectors; v1.1.0^48^) to each component and selecting the one with the lowest inter-peak temporal variance, followed by manual verification.

The data were segmented into epochs between −1 and 1.5 seconds relative to onset of the RDK (visual epochs) and to the end of the reaching movement (motor epochs). Analysis focused on data from 11 sensors above the left sensorimotor area, contralateral to the hand used for movement (i.e., MLC21, MLC22, MLC23, MLC24, MLC31, MLC41, MLC51, MLC52, MLC53, MLC61, MLC62)

### Beta power and burst extraction

We implemented the burst detection algorithm described in Szul et al. ^8^, which iteratively analyzes single trial time-frequency (TF) decompositions until no further bursts are detected above the noise floor. To limit the influence of slower event-related field (ERF) dynamics on burst waveforms, we first averaged the epochs in the temporal domain to compute the ERF and then regressed this out of the signal for each trial. To compute the single-trial TF decompositions, we used the superlet transform with an adaptive order ranging from 1 to 40, a base cycle parameter of 4 ^49^. This TF decomposition was used to compute the power spectral density (PSD) for each sensor by averaging single-trial TF power over time to obtain single-trial PSDs, which were then averaged across trials within each experimental block. The resulting PSDs were parameterized using the specparam algorithm ^23^, estimating the aperiodic (1/f-like) component as a straight line in log-log space. This estimated aperiodic spectrum was subsequently subtracted from each single-trial TF decomposition, and the iterative burst detection algorithm was applied to the residual amplitude, to reduce the influence of broadband power fluctuations and ensure that detected bursts reflect genuine ones^50^.

During each iteration, the algorithm identifies the global maximum amplitude in the TF space, fits a two-dimensional Gaussian to this peak by calculating the symmetric full-width at half maximum (FWHM) in the time and frequency dimensions. This 2D Gaussian parameterization defines burst features in the TF space: peak time, duration, peak amplitude, peak frequency, and frequency span. The Gaussian is then subtracted from the TF decomposition, and the next iteration operates on the resulting residual TF matrix. This process continues until no global maxima above the noise floor remains (2 standard deviations above the mean amplitude over all time and frequency bins, recomputed each iteration). To avoid edge effects near the beta band limits, we applied the algorithm to TF data between 10 and 33 Hz, but only retained bursts with a peak frequency within the beta band (13-30 Hz) for further analysis, following a previously published procedure ^8^.

Using the peak time of each detected burst, we extracted the waveform from the “raw” time series, which was unfiltered except for the 120 Hz low-pass filter applied during preprocessing. To determine the appropriate time window for waveform extraction, we calculated lagged coherence across all trials and sensors from 5 to 40 Hz and 2.5-5 cycles ^51^. We used overlapping time windows with lag- and frequency-dependent widths, and Fourier coefficients were obtained for each time window using a Hann-windowed Fourier transform. The time window for waveform extraction was defined as the full-width at half-maximum (FWHM) of lagged coherence averaged within the beta band, resulting in 5.5 cycles or 255.81 ms at the mean beta frequency of 21.5 Hz, which we rounded up to 260 ms. The time series in this 260 ms window centered on the peak time was then extracted from the trial time series.

To align burst waveforms with the signal deflection corresponding to the peak amplitude, we band-pass filtered them within their detected frequency span (using a zero-phase FIR filter with a Hamming window), computed their instantaneous phase with the Hilbert transform, and re-centered the “raw” (pre-band-pass filtering) waveform around the phase minimum (corresponding to the peak or trough) closest to the peak time detected in the TF space^52^. Bursts were discarded if this time point was more than 30 ms away from the TF-detected peak time. The DC offset was then subtracted from the resulting waveform. Finally, due to uncertainty in the orientation and source location of the dipoles generating the measured sensor signals, we reversed the sign of burst waveforms with a central deflection that was positive^53^. The open-source code for the burst detection algorithm is available at https://github.com/danclab/burst_detection (v0.1), and further details are provided in the associated study ^8^.

### Burst analysis

To classify burst waveform shapes, we applied principal component analysis (PCA) using 20 components implemented in the scikit-learn library^47^. This analysis was performed on aligned waveforms, treating each waveform time point as a feature. The principal components (PCs) were computed from a subset of the data, consisting of 20% of the waveforms, sampled evenly from each subject, block, and epoch, after excluding waveforms outside the 10th-90th percentile of median amplitude (N = 233,921). All detected bursts (N = 14,328,947) were then projected onto each PC, assigning each burst a score for each component that represents the shape of its waveform along that dimension. To determine how many components were meaningful beyond expected noisy fluctuations of the neural field potential, we used a permutation approach^54^: on each iteration, the matrix of waveforms used for PCA was shuffled by column (time point) to remove correlation between features, and PCA was then applied to the shuffled matrix using the same parameters as for the unshuffled subset. The p-value for each PC was determined by the proportion of shuffled PCAs where the variance explained for that PC was lower than for the PCA of the unshuffled data. A lower proportion indicates that the original component was mostly driven by noise. For each component, 100 permutations were run, using an alpha threshold of p = 0.0025 (Bonferroni-corrected for 20 components).

We then selected PCs that define dimensions along which the mean burst waveform shape varied across the epochs compared to a pre visual stimulus baseline. For each PC, the mean burst waveform score was calculated at each time point of the epochs, and the variance of the mean score over the time course of each epoch was used for PC selection. Components with differences in the temporal variance of the mean score between epochs and baseline above a threshold of 0.002 were chosen for further analysis, namely PC7, PC8, PC9 and PC10. Although these PCs define a continuous waveform-shape space, we subdivided the distribution of burst scores along each selected dimension into quartiles. This step was performed to classify bursts into discrete types with systematically different waveform characteristics, enabling comparisons of burst-rate dynamics across task conditions ^8,16^.

### Statistics

#### Behavior

Behavioral statistical analysis was done using linear mixed models using R (version 3.6.1) (R Core Team, 2022) and lme4 (version 1.1.29;^55^). Fixed effects were assessed using Type III Wald χ2 tests (car version 3.1.0;^56^). Pairwise Tukey-corrected follow-up tests were run using estimated marginal means from the *emmeans* package (version 1.7.3;^57^). To assess whether the experimental manipulation had any effects on behavior, we analyzed RTs and error measures using a simple model with block type (baseline, adaptation, washout) and group (implicit, explicit) as fixed effects, and subject-specific intercepts as random effects (as for all models below). We then focused the analysis on adaptation trials using models with coherence (zero, low, medium, high), group, and blocks (continuous variable from 1 to 6, corresponding to the adaptation blocks) as fixed effects.

#### Burst features, burst count and PC-quartile time series

To investigate the relationship between neural activity and behavior, we analyzed trial-wise neural time series, including beta power, overall burst rate, and PC-quartile-specific burst counts, in relation to behavioral measures using linear mixed-effects modeling combined with permutation-based cluster inference. For each neural measure, behavior was modeled as:

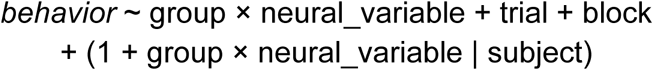

where *neural_variable* corresponded to beta power, overall burst rate, or burst count within a given PC-quartile combination. Behavioral variables of interest included response time and absolute error measures. Group (implicit, explicit) was treated as a fixed effect, trial number and block were included as covariates, and subject-specific random intercepts and slopes were modeled. Separate models were fit for analyses predicting behavior on the current trial and on the subsequent trial.

To identify time periods during which neural activity was significantly related to behavior, we first performed cluster-based permutation tests across time using linear mixed-effects models, as implemented in the clusterperm.lmer function. Permutation tests (10000 iterations) were used to assess the significance of main effects of neural activity as well as group × neural activity interactions over time. Significant clusters were defined as contiguous time points exceeding a cluster-level threshold of *p* < 0.05 and containing at least 15 consecutive samples.

For each significant temporal cluster identified in the permutation analysis, we conducted follow-up window-based linear mixed-effects models by averaging the neural measure across the corresponding time window. These window-averaged neural measures were entered into linear mixed-effects models with the same fixed and random effects structure as above. Type III Wald χ² tests were used to assess the significance of fixed effects. When a significant group × neural activity interaction was observed, post hoc analyses were performed using estimated marginal trends to assess neural-behavior relationships separately within each group, as well as direct contrasts between groups.

## Supplementary Material

**Figure S1.**
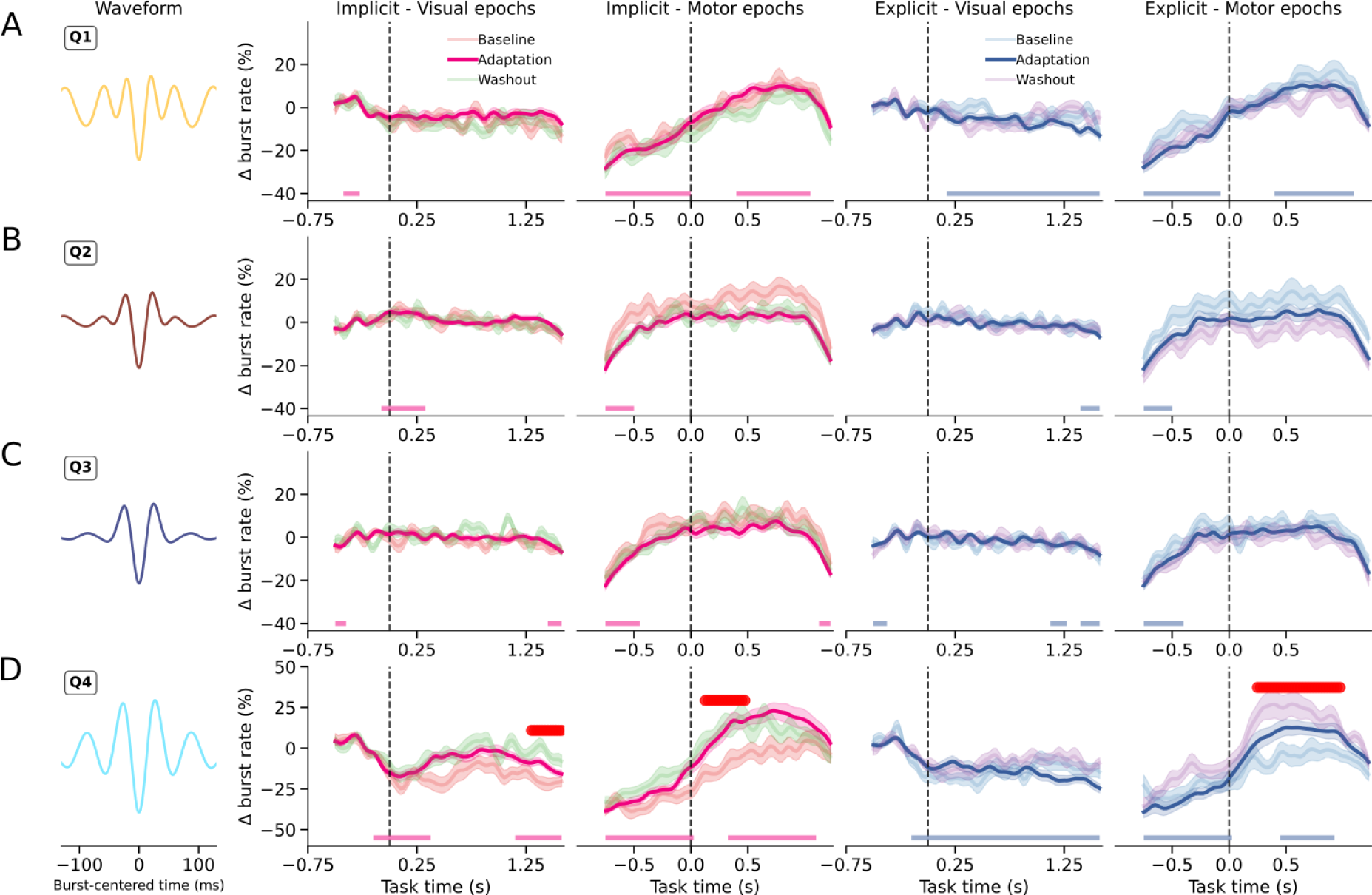
PC7. Baseline-corrected burst rates (Δ burst rate, %) separated by PC7 quartile (Q1-Q4), shown across visual and motor epochs, groups (implicit, explicit), and experimental blocks (baseline, adaptation, washout). Bottom markers indicate time periods where burst rates during adaptation significantly differed from zero. Red markers indicate significant condition differences. Shaded areas represent ± SEM.

**Figure S2.**
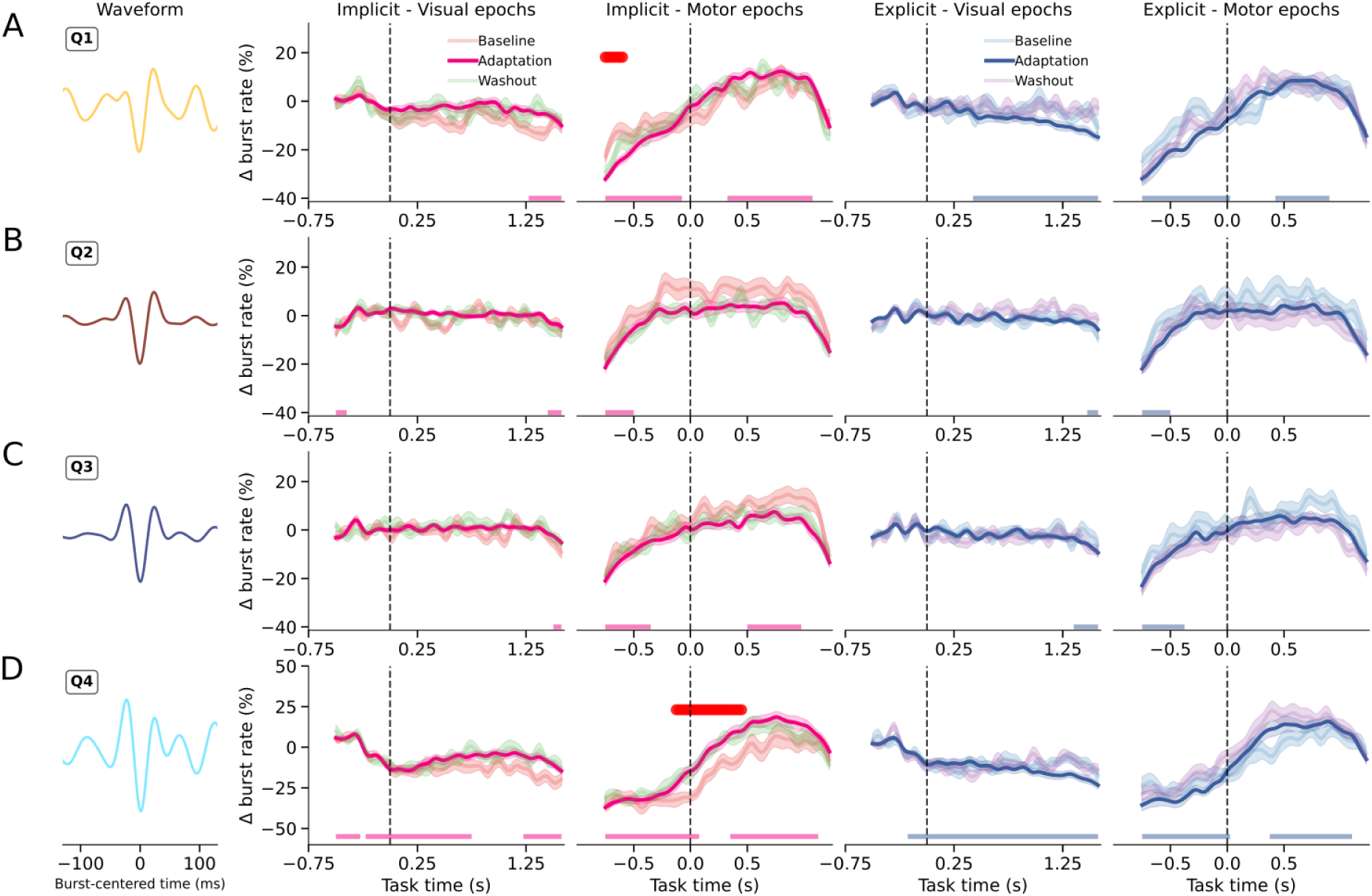
PC9. Baseline-corrected burst rates (Δ burst rate, %) separated by PC7 quartile (Q1-Q4), shown across visual and motor epochs, groups (implicit, explicit), and experimental blocks (baseline, adaptation, washout). Bottom markers indicate time periods where burst rates during adaptation significantly differed from zero. Red markers indicate significant condition differences. Shaded areas represent ± SEM.

**Figure S3.**
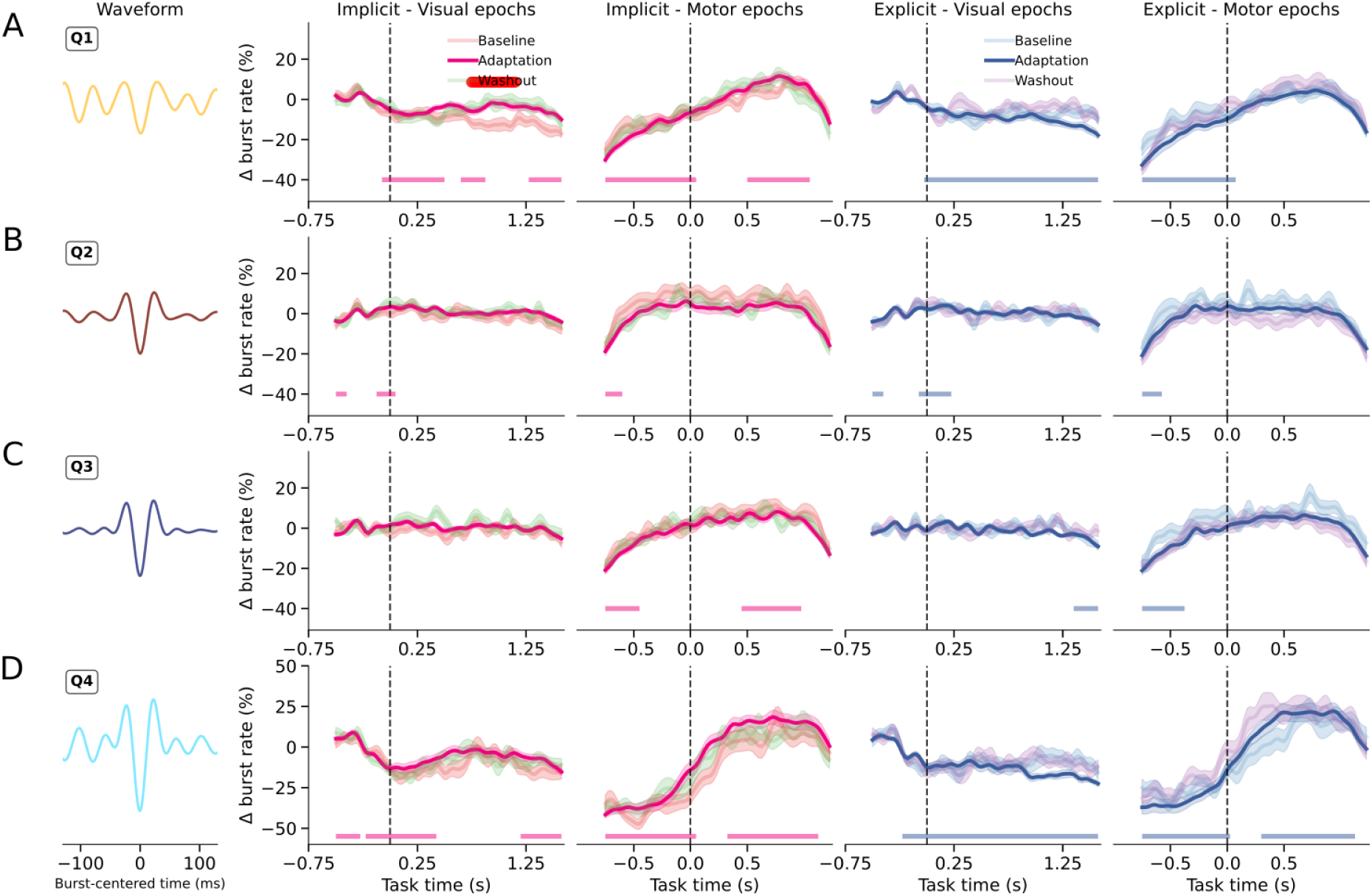
PC10. Baseline-corrected burst rates (Δ burst rate, %) separated by PC7 quartile (Q1-Q4), shown across visual and motor epochs, groups (implicit, explicit), and experimental blocks (baseline, adaptation, washout). Bottom markers indicate time periods where burst rates during adaptation significantly differed from zero. Red markers indicate significant condition differences. Shaded areas represent ± SEM.

## Data and materials availability

All data and code used in this study are available at https://github.com/danclab/explicit_implicit_bursts

